# Visual Experience is not Necessary for the Development of Face Selectivity in the Lateral Fusiform Gyrus

**DOI:** 10.1101/2020.02.25.964890

**Authors:** N. Apurva Ratan Murty, Santani Teng, David Beeler, Anna Mynick, Aude Oliva, Nancy Kanwisher

## Abstract

Here we show robust face-selectivity in the lateral fusiform gyrus of congenitally blind participants during haptic exploration of 3D-printed stimuli, indicating that neither visual experience, nor fovea-biased input, nor visual expertise is necessary for face-selectivity to arise in its characteristic location. Similar resting fMRI correlation fingerprints in individual blind and sighted participants suggest a role for long-range connectivity in the specification of the cortical locus of face-selectivity.

## Introduction

Neuroimaging research over the last 20 years has provided a detailed picture of the functional organization of the cortex in humans. Dozens of distinct cortical regions are each found in approximately the same location in essentially every typically developing adult. How is this complex and systematic organization get constructed over development, and what is the role of experience? Here we address one facet of this longstanding question by testing whether the fusiform face area (FFA), a key cortical locus of the human face processing system, arises in humans who have never seen faces.

Both common sense, and some data, suggest a role for visual experience in the development of face perception. First, faces constitute a substantial percent of all visual experience in early infancy (Smith et al., 2018), and it would be surprising if this rich teaching signal were not exploited. Second, face perception abilities and face-specific neural representations continue to mature for many years after birth (Aylward et al., 2005; Carey et al., 1980; Cohen et al., 2019; Diamond and Carey, 1977; Golarai et al., 2010). Although these later changes could in principle reflect either experience or biological maturation or both, some evidence indicates that the amount (Balas et al., 2019, 2018) and kind (Gilchrist and McKone, 2003; McKone and Boyer, 2006) of face experience during childhood affects face perception abilities in adulthood. In all of these cases, however, it is not clear whether visual experience plays an instructive role in wiring up or refining the circuits for face perception, or a permissive role in maintaining those circuits (Crair, 1999).

Indeed several lines of evidence suggest that some aspects of face perception may develop with little or no visual experience. Within a few minutes, or perhaps even before birth (Reid et al., 2017), infants track schematic faces more than scrambled faces (Johnson, 2012; Johnson et al., 1991). Within a few days of birth, infants can behaviorally discriminate individual faces, across changes in viewpoint and specifically for upright, not inverted faces (Turati, 2004; Turati et al., 2010, 2008). EEG data from infants one to four days old show stronger cortical responses to upright than inverted schematic faces (Buiatti et al., 2019). Finally, functional MRI (fMRI) data show a recognizably adultlike spatial organization of face responses in the cortex of infant monkeys (Livingstone et al., 2017) and 6 month old human infants (Deen et al., 2017). These findings show that many behavioral and neural signatures of the adult face processing system can be observed very early in development, often before extensive visual experience with faces.

To more powerfully address the causal role of visual experience in the development of face processing mechanisms, what is needed is a comparison of those mechanisms in individuals with and without the relevant visual experience. Two recent studies have done just that. Arcaro et al (2017) raised baby monkeys without exposing them to faces (while supplementing visual and social experience with other stimuli), and found that these face-deprived monkeys did not develop face-selective regions of cortex. This finding seems to provide definitive evidence that seeing faces is necessary for the formation of face-selective cortex, as the authors concluded. However, another recent study in humans (van den Hurk et al., 2017) argued for the opposite conclusion. Building upon a large earlier literature providing evidence for category selective responses for scenes (Wolbers et al., 2011), objects, and tools (Amedi et al., 2007; He et al., 2013; Mahon et al., 2009, 2003; Peelen and Downing, 2017; Pietrini et al., 2004) in the ventral visual pathway of congenitally blind participants, the study reported preferential responses for face-related sounds in the fusiform gyrus of congenitally blind humans. These two studies differ in stimulus modality, species, and the nature of the deprivation, and hence their findings are not strictly inconsistent. Nonetheless, they suggest different conclusions about the role of visual experience with faces in the development of face-selective cortex, leaving this important question unresolved.

If indeed true face-selectivity can be found in congenitally blind participants, in the same region of the lateral fusiform gyrus as sighted participants, this would conclusively demonstrate that visual experience is not necessary for face-selectivity to arise in this location. Although the van den Hurk study (2017) provides evidence for this hypothesis, it did not show face-selectivity in individual blind participants, as needed to precisely characterize the location of the activation, and it did not provide an independent measure of the response profile of this region, as needed to establish true face-selectivity (i.e. substantially and significantly higher response to faces than to each of the other conditions tested). Further, the auditory stimuli used by van den Hurk (2017) do not enable the discovery of any selective responses that may be based on amodal shape information (Amedi et al., 2017; Mahon et al., 2009; Striem-Amit et al., 2011) which would be carried by visual or tactile but not auditory stimuli. Here, we use tactile stimuli and individual-subject analysis methods in an effort to determine whether true face-selectivity can arise in congenitally blind individuals with no visual experience with faces.

We also address the related question: Why does the FFA develop so systematically in its characteristic location, on the lateral side of the mid-fusiform sulcus (Weiner et al., 2014)? According to one hypothesis (Amedi et al., 2001; Kanwisher, 2001) this region becomes tuned to faces because it receives preferential input from foveal retinotopic cortex, which in turn disproportionately receives face input (because faces are typically foveated). Another (non-exclusive) hypothesis holds that this region of cortex may have pre-existing feature biases, for example for curved stimuli, leading face stimuli to preferentially engage, and experientially modify, responses in this region (Arcaro and Livingstone, 2017; Hasson et al., 2002; Levy et al., 2001; Malach et al., 2002; Op de Beeck et al., 2019; Powell et al., 2018; Rajimehr et al., 2011; Yue et al., 2014). A third class of hypotheses argue that it is not bottom-up input, but rather interactions with higher-level regions engaged in social cognition and reward, that bias this region to become face-selective (Op de Beeck et al., 2019; Powell et al., 2018). This idea dovetails with the view that category-selective regions in the ventral visual pathway are not just visual processors extracting information about different categories in similar ways, but that each is optimized to provide a different kind of representation tailored to the distinctive post-perceptual use of this information (Peelen and Downing, 2017). Such rich interactions with post-perceptual processing may enable these typically visual regions to take on higher-level functions in blind people (Bedny, 2017; Kim et al., 2017). These three hypotheses make different predictions about face-selectivity in the fusiform gyrus of congenitally blind people, which we test here.

## Results

### Face-selectivity in sighted controls

To validate our methods we first tested for face-selectivity in sighted control participants (N = 15) by scanning them with fMRI as they viewed rendered videos of 3D-printed face, maze, hand and chair stimuli, and as they haptically explored the same stimuli with their eyes closed (Figure 1).

**Figure 1.**
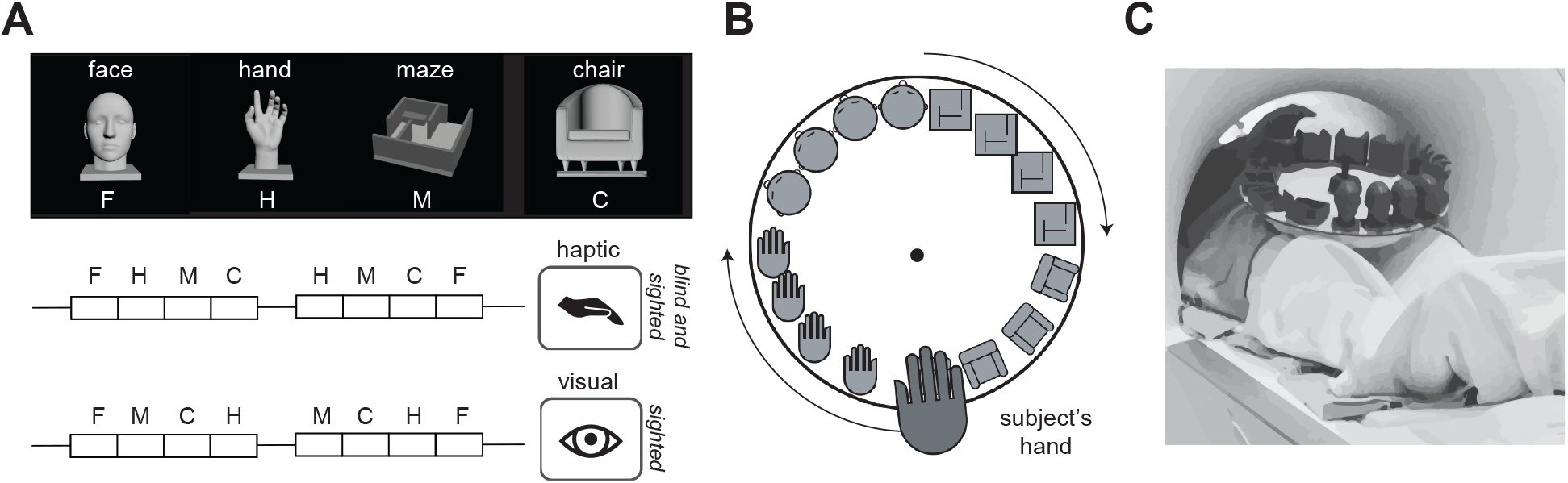
Haptic experiment stimuli and design. **A.** (top) Images showing a rendered example stimulus from each of the 4 stimulus categories used in Experiment 1 - faces (F), hands (H), mazes (M) and chairs (C). (bottom) Experimental design in which the participants haptically explored the stimuli presented in blocks. Sighted participants viewed these rendered stimuli rotating in depth; both sighted and blind subjects haptically explored 3D printed versions of these stimuli. Both haptic and visual experiments included two blocks per stimulus category per run, with 30-second rest blocks at the beginning, middle, and end of each run. During the middle rest period in the haptic condition the turntable was replaced to present new stimuli for the second half of the run. **B.** The stimuli were presented on a rotating turntable, to minimize hand/arm motion. The subjects explored each 3D printed stimulus for 6s after which the turntable was rotated to present the next stimulus. **C.** Image showing an example participant inside the scanner bore extending his hand out to explore the 3D printed stimuli on the turntable.

Whole-brain contrasts of the response during viewing of faces versus viewing hands, chairs, and mazes showed the expected face-selective activations in the canonical location lateral to the mid-fusiform sulcus (Weiner et al., 2014) in both individual participants (Figure 2A), and in a group activation overlap map (Figure 2B). Following established methods in our lab (Cohen et al., 2019; Julian et al., 2012; Norman-Haignere et al., 2016), we identified the top face-selective voxels for each participant within a previously reported anatomical constraint parcel (Julian et al., 2012) for the fusiform face area (FFA), quantified the fMRI response in these voxels in held-out data, and statistically tested selectivity of those response magnitudes across participants (Figure 2C). This method avoids subjectivity in the selection of the functional region of interest (fROI), successfully finds face-selective fROIs in most sighted subjects, and avoids double-dipping or false positives by requiring cross-validation (see Supplemental Figure 3 for control analyses). As expected for sighted participants in the visual condition, the response to faces was significantly higher than each of the other three stimulus categories in held-out data (Figure 2C-D, all *P*<0.005, paired *t*-test).

**Figure 2.**
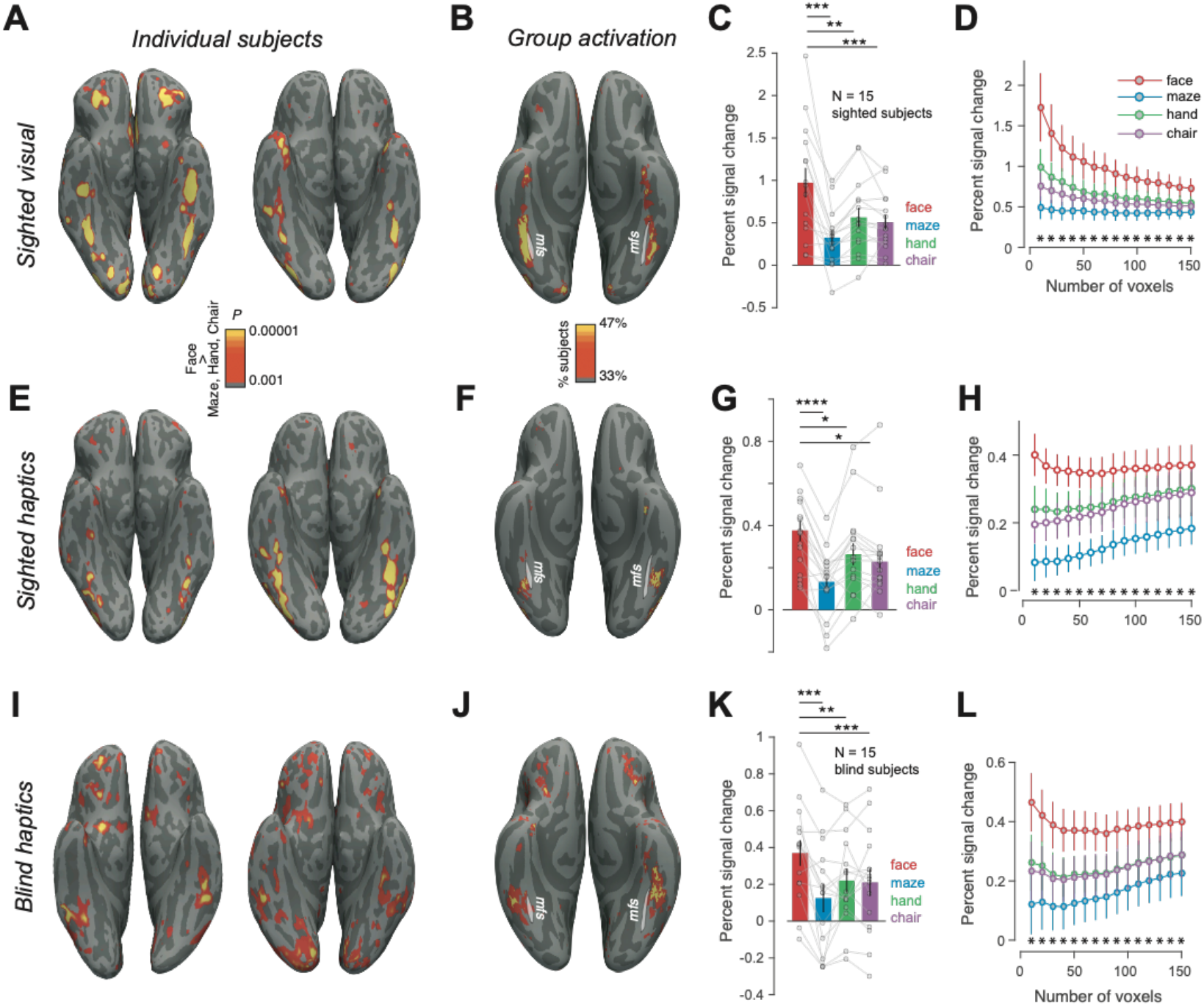
Face selectivity in sighted and blind. **A.** Visual face-selective activations (faces> hands, chairs, and mazes) on the native reconstructed surface for two sighted subjects. **B.** Percent of sighted subjects showing significant visual face selectivity (at the *P*<0.001 uncorrected level in each participant) in each voxel registered to fsaverage surface. White line shows the location of the mid-fusiform sulcus (mfs). **C.** Mean and s.e.m. of fMRI BOLD response across sighted subjects in the top 50 face-selective voxels (identified using independent data) during visual inspection of face, maze, hand and chair images. Individual subject data are overlaid as connected lines. * is *P* < 0.05, ** is P< 0.005 and **** is P < 0.00005 **D.** Response profile of visual face-selective region (in held-out data) in sighted participants as a function of fROI size (error bars indicate the s.e.m.). Stars * indicate significant (P<0.05) difference between faces and each of the 3 other stimulus categories across subjects **E-H.** Same as **A-D** but for sighted subjects during haptic exploration of 3D-printed stimuli. **I-L.** Same as **A-D** but for blind subjects during haptic exploration of 3D-printed stimuli.

In the haptic condition, whole-brain contrasts revealed face-selective activations in a similar location to visual face-selectivity in individual sighted participants (Figure 2E) and in the group overlap map (Figure 2F). fROI analyses using haptic data to both select and quantify held-out responses revealed significantly higher response to faces than each of the three other conditions (all P<0.05, paired *t*-test). Further, when the visual data from the same subject were used to define face-selective fROIs, we observed similar haptic face-selectivity (all P<0.05, paired *t*-test). Note that the absolute magnitude of the fMRI signal was lower for the haptic condition in sighted participants relative to the visual condition, but the selectivity of the response was similar in the two modalities (*t*(28)=0.13, *P*=0.89, unpaired *t*-test, on selectivity index; see Methods for a discussion on the utility of the selectivity index). The observation of haptic face-selectivity in sighted participants demonstrates the effectiveness of our stimuli and methods, and presumably reflects visual imagery of the haptic stimuli (Amedi et al., 2005; O’Craven and Kanwisher, 2000). But the sighted haptic responses do not resolve the main question of this paper, of whether face-selectivity in the fusiform can arise without visual experience with faces. To answer that question, we turn to congenitally blind participants.

### Face-selectivity in blind participants

To test whether blind participants show selectivity for faces despite their lack of visual experience, we scanned congenitally blind participants on the same paradigm as sighted participants, as they haptically explored 3D-printed face, maze, hand and chair stimuli. Indeed, whole-brain contrast maps reveal face-selective activations in the group activation overlap map (Figure 2J) and in most blind subjects analyzed individually (Figure 2I and Supplementary Figure 1). These activations were found in the canonical location (Figure 2I-J), lateral to the mid-fusiform sulcus (Weiner et al., 2014). Following the method used for sighted participants, we identified the top haptic face-selective voxels for each subject within the previously published anatomical constraint parcels for the visual FFA (Supplementary Fig. 2), and measured the fMRI response in these voxels in held-out data. The response to haptic faces was significantly higher than to each of the other three stimulus classes (all *P*<0.005, paired *t*-test). Although the absolute magnitude of the fMRI signal was lower for blind participants haptically exploring the stimuli than for sighted participants viewing the stimuli, the selectivity of the response was similar in the two groups (*t*(28)=0.05, *P*=0.96, unpaired *t*-test, on selectivity index between blind-haptics and sighted-visual and *t*(28)=0.123, *P*=0.90, on selectivity index between blind-haptics and sighted-haptics). Note that fROIs were defined in each participant as the top 50 most significant voxels within the anatomical constraint parcel, even if no voxels actually reached significance in that participant, thus providing an unbiased measure of average face-selectivity of the whole group. These analyses reveal clear face-selectivity in the majority of congenitally blind participants, in a location similar to where it is found in sighted participants. Thus, seeing faces is not necessary for the development of face-selectivity in the lateral fusiform gyrus.

Three further analyses support the similarity in anatomical location of the face-selective responses for blind participants feeling faces and sighted participants viewing faces. First, a whole-brain group analysis found no voxels showing an interaction of subject group (sighted visual versus haptic blind) by face/nonface stimuli even at the very liberal threshold of *P*<0.05 uncorrected, aside from a region in the collateral sulcus that showed greater scene selectivity in sighted than blind participants. Second, a new fROI-based analysis using hand-drawn anatomical constraint region for the FFA based on precise anatomical landmarks from Weiner et al (2014) showed similar face-selectivity in the haptic blind data (see Supplemental Figure 3). Finally, there is no evidence for differential lateralization of face-selectivity in haptic blind versus visual sighted (see Supplemental Figure 4). Taken together, these analyses suggest that haptic face-selectivity in the blind arises in a similar anatomical location to visual face-selectivity in the sighted.

### Why do face-selective activations arise where they do?

Our finding of haptic face-selective activations in the fusiform gyrus of blind subjects raises another fundamental question – why does face-selectivity arise so systematically in that specific patch of cortex, lateral to the mid-fusiform sulcus (Op de Beeck et al., 2019)? According to one widespread hypothesis, this cortical locus becomes face-selective because it receives greater input from foveal (versus peripheral) retinotopic cortex, where face stimuli most often occur (Arcaro and Livingstone, 2017; Hasson et al., 2002; Levy et al., 2001; Malach et al., 2002). This hypothesis cannot account for the face-selectivity in the blind observed here, because these subjects never received any visual input from the fovea at all.

According to another widely-discussed hypothesis, this region is biased for curved stimuli, as part of an early-developing shape-based proto-map upon which higher-level face-selectivity is subsequently constructed (Srihasam et al., 2014). If this proto-map is amodal, responding also to haptic shape (Amedi et al., 2017, 2010), haptic face-selective regions in blind participants should respond preferentially to curved compared to rectilinear shapes. We tested this hypothesis in a second experiment in which seven of our original congenitally blind participants returned for a new scanning session in which they haptically explored a stimulus set comprising the four stimulus categories used previously and two additional categories: spheroids and cuboids. This experimental design enabled us to both replicate face-selectivity in an additional experimental session and determine whether the face-selective voxels were preferentially selective for curvilinear shapes (spheroids) over rectilinear shapes (cuboids). Face-selective activations were replicated in the second experiment session (Figure 2a, top right panel). In a strong test of the replication, we quantified face-selectivity by choosing the top face-selective voxels from the first experiment session within the FFA parcel, spatially registered these data to those from the second session in the same participant, and measured the response of this same fROI in the second experimental session. We replicated the face-selectivity from our first experiment (Fig. 2b). Further, we found that the face-selective voxels did not respond more strongly to spheroids than cuboids (*t*(6)=-0.65, *P*=0.54, paired *t*-test). These data argue against a curvature bias in a pre-existing amodal shape map as a determinant of the cortical locus of face-selectivity in blind subjects.

If it is neither early-developing retinotopic nor feature-based proto-maps (Arcaro and Livingstone, 2017; Srihasam et al., 2012) that specify the cortical locus of face-selectivity in blind participants, what does? In sighted subjects, category-selective regions in the ventral visual pathway serve not just to analyze visual information, but to extract the very different representations that serve as inputs to higher-level regions engaged in social cognition, visually-guided action, and navigation (Kim et al., 2017; Op de Beeck et al., 2019; Peelen and Downing, 2017; Powell et al., 2018). Perhaps it is their interaction with these higher-level cortical regions that drives the development of face-selectivity in this location (Op de Beeck et al., 2019; Powell et al., 2018). The previously mentioned preferential response to face-related sounds in the fusiform of blind subjects (van den Hurk et al., 2017) provides some evidence for this hypothesis. To test the robustness of that result, and to ask whether face-selectivity in blind participants arises in the same location for auditory and haptic stimuli we next ran a close replication of the Van den Hurk et al (2017) study on seven of the congenitally blind subjects in our pool.

Participants heard the same short clips of face, body, object and scene related sounds used in the previous study (van den Hurk et al., 2017), while being scanned with fMRI. Examples of face-related sounds included audio clips of people laughing or chewing, body-related sounds included clapping or walking, object-related sounds included a ball bouncing and a car starting, and scene-related sounds included clips of waves crashing and a crowded restaurant. Indeed, we found robustly selective responses to face sounds compared to object, scene and body-related sounds in individual blind participants, replicating van den Hurk (2017). Auditory face activations were found in similar locations as haptic face activations (Fig 2a and Supplementary Fig. 2). To quantify auditory face-selectivity we chose face-selective voxels in the FFA parcels from the haptic paradigm and tested them on the auditory paradigm. This analysis revealed clear selectivity for face sounds across the group (Fig. 2c) and further showed that auditory face-selectivity co-localizes with haptic face-selectivity in blind participants.

### Do Similar Connectivity Fingerprints Predict the Locus of Face-Selectivity in the Sighted and Blind?

The question remains: What is it about the lateral fusiform gyrus that marks this region as the locus where face-selectivity will develop? A longstanding hypothesis holds that the long-range connectivity of the brain, much of it present at birth, constrains the functional development of cortex (Bi et al., 2016; Dubois et al., 2014; Op de Beeck et al., 2019; Osher et al., 2016; Peelen et al., 2017; Striem-Amit et al., 2015; Sur et al., 1986; Wang et al., 2017, 2015a). Evidence for this hypothesis comes from demonstrations of distinctive ‘connectivity fingerprints’ of many cortical regions in adults (Osher et al., 2016; Passingham et al., 2002; Saygin et al., 2012), including the fusiform face area (Osher et al., 2016; Saygin et al., 2012). To test this hypothesis in our group of sighted and blind participants, we used resting-state fMRI correlations as a proxy for long-range connectivity (Osher et al., 2019). We first asked whether the ‘correlation fingerprint’ of face-selective regions was similar in blind and sighted. The fingerprint of every vertex in the fusiform was defined as the correlation between the resting-state time-course of that vertex and the average time-course within each of 355 cortical regions from a standard whole-brain parcellation (Glasser et al., 2016). Fingerprints corresponding to face-selective vertices were highly correlated between sighted and blind subjects in both hemispheres. Specifically, the 355-dimensional vector of correlations between the face-selective voxels (averaged across the top 200 most face-selective voxels within each hemisphere and individual, then averaged across participants) and each of the 355 anatomical parcels was highly correlated between the blind and sighted participants (mean±std across 1000 bootstrap estimates across subjects, left hemisphere Pearson R = 0.82±0.08, right hemisphere Pearson R = 0.76±0.09). In contrast, the analogous correlations between the with best face-selective and scene-selective vertices, bootstrap resampled 1000 times across subjects, left hemisphere Pearson R = 0.55±0.12, right hemisphere Pearson R = 0.60±0.09, *P*=0, permutation test between face and scene selective on Fisher transformed correlations)

Next, we tested a stronger version of the connectivity hypothesis by building computational models that learn the mapping from the correlation fingerprint of each vertex to the functional activations. Specifically, following established methods (Osher et al., 2019; Saygin et al., 2012, we trained models to learn the voxel-wise relationship between correlation fingerprints and face-selectivity, and tested them using leave-one-subject-out cross-validation. We first tested the efficacy of this approach within the sighted and blind groups separately. Models trained on data within each group (blind or sighted) on average predicted the spatial pattern of each held-out subject’s face-selective activations significantly better than a group analysis of the functional selectivity from the other participants in that group (Fig. 3b, *P*<0.05, paired *t*-test with predictions from a random-effects group analysis). This result shows that it is indeed possible to learn a robust mapping from voxel-wise correlation fingerprints to voxel-wise face-selective functional activations within each group. We then asked whether the model trained on one group of subjects (e.g., sighted participants), would generalize to the other group of subjects (e.g., blind participants), as a stronger test of the similarity in the connectivity of face-selective regions between sighted and blind participants. Indeed, these predictions of the spatial pattern of face-selectivity were found not only for the held-out subjects within each group, but also for subjects in the other group, significantly outperforming the predictions from the functional activations of other participants within the same group (Fig. 3b, *P*<0.05, paired *t*-test with predictions from a random-effects group analysis). Although resting functional MRI correlations are an imperfect proxy for structural connectivity (Buckner et al., 2013), these results suggest that face-selectivity in the fusiform is predicted by similar long-range connectivity in sighted and blind participants.

**Figure 3.**
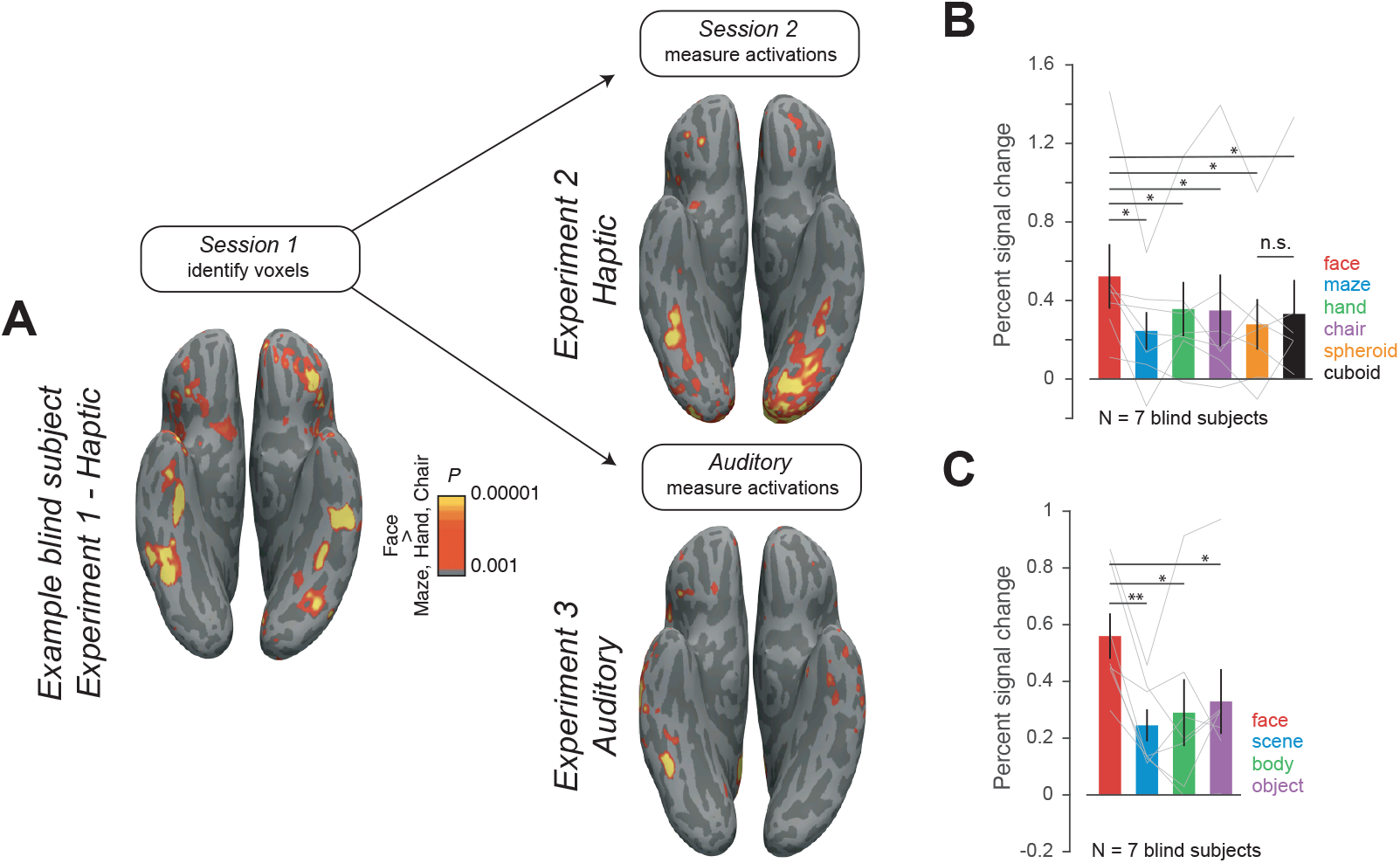
Responses of face-selective regions in blind participants during haptic exploration of curved stimuli and auditory listening to sound categories. **A.** Face selective activation (faces > hands, chairs, and scenes) in an example blind subject during haptic exploration of 3D printed stimuli in Session 1 (left), replicated in an additional follow-up scan with additional curved stimuli (right top) and in an auditory paradigm (bottom right). Right: Mean and s.e.m. across participants of fMRI BOLD response in the top 50 face-selective voxels (identified from Session 1) during **B.** haptic exploration of spheroids and cuboids (in addition to face, maze, hand, chair stimuli as before) and **C.** auditory presentation of face, scene, body and object related sounds. In **B** and **C**, individual subject data are overlaid as connected lines; Lines above the bars indicate all Ps < 0.05 (two-tailed t-test). n.s. is not significant.

**Figure 4.**
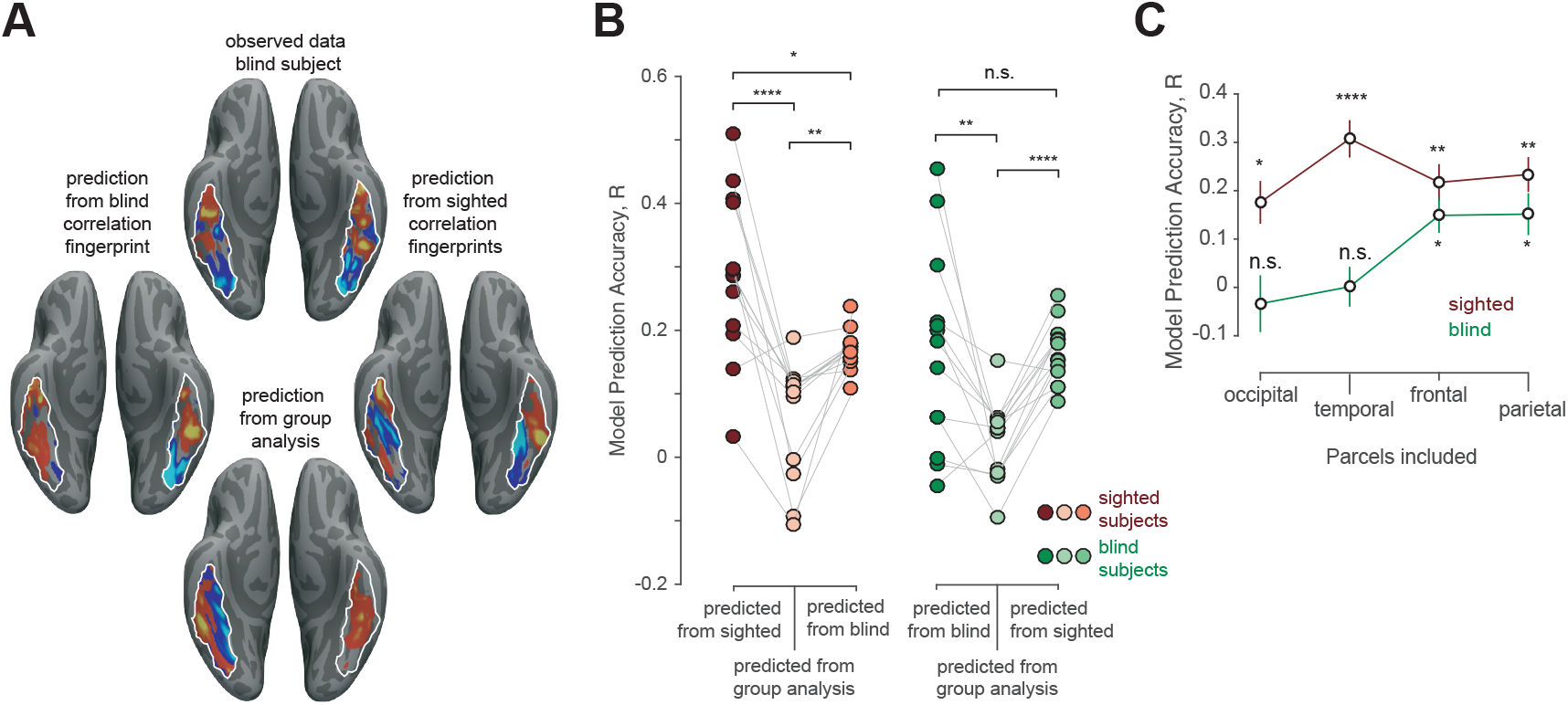
“Correlation fingerprints” predict the spatial locations of face selectivity. **A.** Top: Observed face-selective activation in an example blind subject (z-scored units) Predicted activations for the same subject based on that individual’s correlation fingerprint (CF) using a model trained on other blind subjects’ haptic activations (left), or on sighted subjects’ visual activations (right). Bottom: predicted activation based on group analysis of face-selective responses in other blind subjects. **B.** Model prediction performance across blind and sighted subjects. Each dot is the model prediction accuracy for a single sighted (left, red) and blind (right, green) subject. Model predictions were obtained from CFs derived from either the same group (left column in each set), the functional group analysis of the remaining subjects (middle column in each set) or from CFs of the opposite group (right column in each set). Paired *t*-tests were performed on Fisher-transformed data. * is *P*<0.05, ** is *P*<0.005, **** is *P*<0.00005 and n.s. is not significant. **C.** Model prediction accuracy of face selectivity for sighted (visual responses) and blind (haptic responses) based on CFs including only target regions within one lobe at a time. Statistics indicate significantly better prediction from CFs than from the random effects analysis of face selectivity in the rest of the group; calculations and notations same as **B.**

The previous analysis does not tell us *which* connections are most predictive of the functional activations in sighted and blind participants. In order to address this question, we re-analyzed the correlation fingerprints using only target regions from a single lobe (frontal, temporal, occipital and temporal) at a time. Here we find that although voxel-wise visual face-selectivity in sighted subjects was significantly predicted from correlation fingerprints to each of the four lobes of the brain individually (*P*<0.05), in blind subjects, haptic face-selectivity was significantly predicted from parietal and frontal regions only. These findings implicate top-down inputs in the development of face-selectivity in the fusiform in blind subjects.

## Discussion

How are functionally specific cortical regions wired up in development, and what is the role of experience? This study tests three widely discussed (non-exclusive) hypotheses about the origins of the fusiform face area (FFA), and presents evidence against each of them. Our finding of robust face-selective responses in the lateral fusiform gyrus of most congenitally blind participants indicates that neither foveal input, nor perceptual expertise, nor indeed any visual experience at all is necessary for the development of face-selective responses in this region. Our further finding that the same region does not respond more to curvy than rectilinear shapes in blind participants casts doubt on the hypothesis that face-selectivity arises where it does in the cortex because this region constitutes a curvature-biased part of an early-developing proto-map. Finally, our finding of very similar correlation fingerprints predictive of face-selectivity for sighted and blind participants, particularly from parietal and frontal regions, supports the hypothesis that top-down connections play a role in specifying the locus where face-selectivity develops.

Our findings extend previous work in two important ways. First, although previous studies have reported some evidence for cortical selectivity in congenitally blind participants for stimulus categories such as scenes and large objects (He et al., 2013; Wolbers et al., 2011), tools (Peelen et al., 2013) and bodies (Kitada et al., 2014; Striem-Amit and Amedi, 2014), prior work has either failed to find face-selectivity in the fusiform at all in blind participants (Goyal et al., 2006; Kitada et al., 2013; Pietrini et al., 2004) or reported only preferential responses in a group analysis (van den Hurk et al., 2017). Our analyses meet a higher bar for demonstrating face-selectivity by showing i) robust face-selectivity in *individual* blind subjects, ii) significantly stronger responses of these regions to *each* of three other stimulus conditions, iii) a similar selectivity index of this region for blind and sighted participants, iv) internal replications of face-selectivity in the fusiform across imaging sessions and across sensory modalities, and v) a demonstration that haptic face-selectivity in the blind cannot be explained by selectivity for haptic curvature. Second, our findings provide evidence against the most widespread theories for why face-selectivity develops in its particular stereotyped cortical location. Apparently, neither foveal face experience, nor a bias for curvature is necessary. Instead, we present evidence that the pattern of structural connections to other cortical regions may play a key role in determining the location where face-selectivity will arise. In particular, we demonstrated that the correlation fingerprint predictive of visual face-selectivity for sighted subjects can also predict the pattern of haptic face-selectivity in blind subjects (and vice versa). Using this predictive modeling approach we also find preliminary evidence for a dominant role of top-down connections from frontal and parietal cortices in determining the locus of face-selectivity in blind subjects (Powell et al., 2018).

Many important questions remain. First, is face-selectivity innate, in the sense of requiring no perceptual experience with faces at all? The present study does not answer this question because even though our congenitally blind participants have never seen faces, they have certainly felt them and heard the sounds they produce. For example blind people do not routinely touch faces during social interactions, but certainly feel their own face and occasionally the faces of loved ones. It remains unknown whether this haptic experience with faces is necessary for the formation of face-selectivity in the fusiform.

Second, what is actually computed in face-selective cortex in the fusiform of blind participants? It seems unlikely that the main function of this region in blind people is haptic discrimination of faces, a task they do rarely. One possibility is that the “native” function of the FFA is individual recognition, which is primarily visual in sighted people (i.e., face recognition) but can take on analogous functions in other modalities in the blind (Amedi et al., 2017; Bi et al., 2016; Mahon et al., 2009; Merabet et al., 2005; Wang et al., 2015b). such as voice identity processing (Hölig et al., 2014b, 2014a). This hypothesis has received mixed support in the past (Dormal et al., 2017; Fairhall et al., 2017) but could potentially account for the observed higher responses to haptic faces and auditory face-related sounds found here. Another possibility is that this region takes on a higher-level social function in blind people, such as inferring mental states in others (but see Bedny et al., 2009). An interesting parallel case is the finding that the “visual word form area” in the blind actually processes not orthographic but high-level linguistic information (Kim et al., 2017). Similarly if the “blind FFA” in fact computes higher-level social information, that could account for another puzzle in our data, which is that not all blind participants show the face-selective response to haptic and auditory stimuli. This variability in our data is not explained by variability in age (correlation of age with selectivity index = 0.03, p=0.89), sex (*R*= −0.12, *P*=0.65), or task performance (*R*= 0.06, *P*=0.81). Perhaps the higher-level function implemented in this region in blind participants is not as automatic as visual face recognition in sighted.

Finally, while our results are clearly consistent with those of van den Hurk et al (2017), they seem harder to reconcile with the prior finding that monkeys reared without face experience do not show face-selective responses (Arcaro et al., 2017). Importantly, though, auditory and haptic responses to faces have not to our knowledge been tested in face-deprived monkeys, and it is possible they would show the same thing we see in humans. If so it would be informative to test whether auditory or tactile experience with faces during development is necessary for face-selective responses to arise in monkeys deprived of visual experience with faces (something we cannot test in humans). In advance of those data, a key difference to note between the face deprivation findings in monkeys and the blind data from humans is that face-deprived monkeys confronting visual faces for the first time presumably had no idea what those visual patterns meant, whereas all the blind participants in our study immediately recognized that the 3D-printed stimuli were faces, without being told. Detecting the presence of a fellow primate may be critical for obtaining a face-selective response.

In sum, we show that visual experience is not necessary for face-selectivity to develop in the lateral fusiform gyrus, and neither apparently is a feature-based proto-map in this region of cortex. Instead, our data suggest that the long-range connectivity of this region, which develops independent of visual experience, may mark the lateral fusiform gyrus as the site of face-selective cortex.

## Methods

### Participants

15 sighted and 15 blind subjects participated in the experiment (6 females in the sighted and 7 females in the blind group, mean ± s.e.m. age 29 ± 2 years for sighted; 28 ± 2 years for blind participants). Two additional blind subjects who were recruited had to be excluded because they were not comfortable in the scanner. Seven subjects from the blind pool returned participated in Experiments 2 and 3 in an additional experiment session (4 females, mean ± s.e.m. age 28 ± 3y). All studies were approved by the Committee on the Use of Humans as Experimental subjects of the Massachusetts Institute of Technology (MIT). Participants provided informed written consent before the experiment and were compensated for their time. All blind participants recruited for the study were either totally blind or had only light perception from birth. None of our subjects reported any memory of spatial or object vision. Details on the blind subjects are summarized in Supplemental Table 1.

### Stimuli

*Experiment 1.* In Experiment 1, blind subjects explored 3D-printed stimuli haptically and sighted subjects explored the same stimuli visually and haptically. The haptic stimuli comprised 5 exemplars each from 4 stimulus categories – faces, mazes, chairs and hands. The 3D models for the face stimuli were generated using FaceGen 3D print (software purchased and downloaded from facegen.com/3dprint.htm). Face stimuli were printed on a 4×4cm base and were 7cm high. 3D models of mazes were similar to house layouts and were designed on Autodesk 3ds Max (Academic License, v2017). The maze layouts were 5×5 cm and consisted of straight walls with entryways and exits and a small raised platform. The 3D models for the hand stimuli were purchased from Dosch Design (Dosch 3D: Hands dataset, from doschdesign.com). The downloaded 3D stimuli were thereafter customized to remove the forearm and include only the wrist and the hand. The stimuli were fixed to a 4×4cm base and were ∼7cm high and included 5 exemplar hand configurations. Finally, the 3D models for chairs were downloaded from publicly available databases (from archive3d.net). The stimuli were all armchairs of different types and were each printed on a 5×5cm base and were ∼7cm tall. Because subjects performed a one-back identification task inside the scanner, we 3D-printed 2 identical copies of each stimulus, generating a total set of 40 3D-printed models (4 object categories x 5 exemplars per category x 2 copies). The 3D models were printed on a FormLabs stereolithography (SLA) style 3D-printer (Form-2) using the grey photopolymer resin (FLGPGR04). This ensured that the surface texture properties could not distinguish the stimulus exemplars or categories. The 3D-prints generated from the SLA style 3D-printers have small deformities at the locations where the support scaffolding attaches with the model. We ensured that these deformities were not diagnostic by either using the same pattern of scaffolding (for the face and hand stimuli) or filing them away after the models were processed and cured (chair stimuli). No deformities were not present on the mazes which could be printed directly without the support scaffolds. Nonetheless, we instructed subjects to ignore small deformities when judging the one-back conditions. Visual stimuli for the sighted subjects included short 6s video animations of the 3D renderings of the same 3D printed stimuli (in grey on a black background) rotating in depth. The animations were rendered directly from Autodesk 3Ds Max. Each stimulus subtended about 8 degrees of visual angle around a centrally located black fixation dot. The .STL files used to 3D print the different stimuli and the animation video files will be made available upon request.

*Experiment 2.* Stimuli for Experiment 2 included stimuli from Experiment 1 (face, maze, hand and chair) and 2 additional object categories: spheroids and cuboids. The 3D models were designed from scratch on Autodesk 3Ds max. The spheroids were egg-shaped and the cuboids were box-shaped and were each printed on a 4×4cm base. We 3D-printed 2 copies each of 4 exemplars each of the spheroids and cuboids (3 variations in the x, y and z plane plus one sphere/cube) that varied in elongation. Each stimulus was ∼7cm high.

*Experiment 3.* The auditory stimuli used in Experiment 3 were downloaded from https://osf.io/s2pa9/, graciously provided by the authors of a previous study(van den Hurk et al., 2017). Each auditory stimulus was a short ∼1800ms audio clip and the stimulus set consisted 64 unique audio clips, 16 each from one of 4 auditory categories (face, body, object and scene). Example of face-related stimuli included recorded sounds of people laughing and chewing, body-related stimuli included audio-clips of people clapping and walking, object-related sounds included sounds of a ball bouncing and a car starting and scene-related sounds included clips of waves crashing and a crowded restaurant. The overall sound intensity was matched across stimuli by normalizing the rms value of the sound pressure levels. We did not perform any additional normalization of the stimuli, except to make sure that the sound intensity levels were within a comfortable auditory range for our subjects.

### Paradigm

All 15 blind and 15 sighted subjects participated in Experiment 1. Blind subjects explored 3D-printed faces, mazes, hands and chairs presented on a MR-compatible turn-table inside the fMRI scanner while performing an orthogonal one-back identity task (mean subject accuracy on the one-back task: 79%, 81%, 89% and 86% for face, maze, hand and chair stimuli respectively). Each run contained three 30-second rest periods at the beginning, middle, and end of the run, and eight 24-second long haptic stimulus blocks (two blocks per stimulus category, see Supplemental Fig 1). During each haptic stimulus block, blind subjects haptically explored four 3D-printed stimuli (3 unique stimuli plus a one-back stimulus) in turn for 6 seconds each, timed by the rotation of the turntables the stimuli were velcroed to (Supplemental Fig. 1). Once every stimulus had been explored across four blocks (one per stimulus category), the turntable was replaced during the middle rest period with a new ordering of stimuli (Supplemental Fig 1). The haptic sessions were interleaved with 3-4 minute resting-state scans during which subjects were instructed to remain as still as possible. Each turntable included 16 stimuli (4 stimulus categories x 4 exemplars per category) and the stimulus and category orders were randomized across experimental runs (by changing the position of the stimuli on the turntable). Each run lasted 4min42s and 14/15 blind subjects completed 10 runs (one subject completed 8 runs) of the experiment. Blind subjects were not trained on the stimuli or the task but were only familiarized with the procedure for less than 5 minutes prior to scanning. Care was also taken to ensure that no information about any of the stimulus categories was provided to the participants prior to scanning. Sighted participants performed the analogous task visually (mean accuracy on the one-back task: 99%, 97%, 99% and 99% for face, maze, hand and chair stimuli respectively), viewing renderings of the same 3D-printed stimuli rotating in depth, presented at the same rate and in the same design (four exemplar stimuli per stimulus category per block). Sighted subjects completed 5 runs over the course of an experimental session.

A subset of blind subjects (N=7) returned for a second session (a few days to a few months after the first session) and participated in Experiments 2 and 3. In Experiment 2, subjects performed 10 runs of a similar haptics paradigm as in Experiment 1, but the turntables included 2 additional stimulus categories beyond faces, mazes, hands and chairs: spheroids and cuboids. In the additional session, runs of Experiment 2 were alternated with runs from Experiment 3, which was a direct replication of the auditory paradigm from a previously published study(van den Hurk et al., 2017). The stimulus presentation and task design etc. were the same as in the previous study(van den Hurk et al., 2017). Briefly, subjects were instructed to compare the conceptual dissimilarity of each auditory stimulus with the preceding auditory stimulus on a scale of 1-4. Each auditory stimulus was a short ∼1800ms audio clip and the stimulus set consisted 64 unique audio clips, 16 each from one of 4 auditory categories (face, body, object and scene). The stimuli were presented in a block design, with 4 blocks per category in each run. Each block consisted eight stimuli per category, chosen at random from the 16 possible stimuli set. Each run lasted 7min30s and each subjects participated in 9 runs of this auditory paradigm interleaved with the Experiment 2 haptic paradigm.

### Data acquisition

All experiments were performed at the Athinoula A. Martinos Imaging Center at MIT on a Siemens 3-T MAGNETOM Prisma Scanner with a 32-channel head coil. We acquired a high-resolution T1-weighted (multi-echo MPRAGE) anatomical scans at the first scan (Acquisition parameters: 176 slices, voxel size: 1 x 1 x 1 mm, repetition time (TR) = 2500ms, echo time (TE) = 2.9ms, flip angle = 8 °). Functional scans included 141 and 225 T2*-weighted echoplanar (EPI) BOLD images for each haptic and auditory run respectively (Acquisition parameters: simultaneous interleaved multi-slice acquisition (SMS) 2, TR=2000ms, TE=30ms, voxel-size 2mm isotrotropic, number of slices=52, flip angle: 90°, echo-spacing 0.54ms, 7/8 phase partial fourier acquisition). Resting state data were acquired for 13/15 blind and 13/15 sighted subjects that participated in Experiment 1 (scanning parameters: TR = 1500ms, TE = 32ms, voxel-size 2.5 x 2.5 x 2.5mm isotropic, flip angle 70°, duration of resting state data: 27.5±0.5 minutes for sighted and 27.2 ± 0.5 min for blind subjects). Sighted participants closed their eyes during the resting state runs.

### Data Preprocessing and Modeling

fMRI data preprocessing and generalized linear modeling were performed on Freesurfer (version: 6.0.0; Downloaded from: https://surfer.nmr.mgh.harvard.edu/fswiki/). Data preprocessing included slice time correction, motion correction of each functional run, alignment to each subject’s anatomical data, and smoothing using a 5mm FWHM Gaussian kernel. Generalized linear modelling included one regressor per stimulus condition, as well as nuisance regressors for linear drift removal and motion correction per run. Data were analyzed in each subjects’ native volume (analysis on the subjects’ native surface also resulted in qualitatively similar results). Activation maps were projected on the native reconstructed surface to aid visualization. For group-level analyses, data were co-registered to standard anatomical surface coordinates using Freesurfer’s *fsaverage* (MNI305) template. Because the exact anatomical location of the FFA varies across participants, this activation often fails to reach significance in a random effects analysis, even with a sizable number of sighted participants. So to visualize the average location of face selective activations in each group, each voxel was color-coded according to the number of participants showing a selective activation in that voxel.

Resting-state data were preprocessed using the CONN toolbox (https://www.nitrc.org/projects/conn). Structural data were segmented and normalized. Functional data were motion-corrected and nuisance regression was performed to remove head-motion artifacts and ventricular and white-matter signals. Subject motion threshold was set at 0.9mm to remove timepoints with high motion. The resting-state data were thereafter filtered with a bandpass filter (0.009 to 0.08 Hz). All data were projected onto the *fsaverage* surface using Freesurfer (<u>freesurfer.net</u>) for further analysis.

### fROI analysis in the fusiform

We used functional region-of-interest (ROI) analyses in sighted and blind subjects to quantify face selectivity. Specifically, we used previously published(Julian et al., 2012) anatomical constraint parcels downloaded from https://web.mit.edu/bcs/nklab/GSS.shtml within which to define the Fusiform Face Area (FFA). These parcels were projected into each subjects’ native volume. The face-selective ROI for each subject was identified as the top *N* voxels within the anatomical constraint parcel for the FFA that responded most significantly to face relative to maze, hand and chair stimuli (regardless of whether any of those voxels actually reached any statistical criterion). We fixed *N* as the 50 top voxels (or 400 mm^3^) prior to analyzing the data, but also varied from 10-150 voxels (i.e., 80 – 1200 mm^3^) to estimate selectivity as a function of ROI size. We always used independent data to select the top voxels and estimate the activations (based on an odd-even run cross-validation routine) to avoid double-dipping. The betas were converted into BOLD percent signal change values by dividing by the mean signal strength. The statistical significance was assessed using two-tailed paired *t*-tests between object categories across subjects.

### Correlational-fingerprint based prediction

This analysis tests how well we could predict the functional activations in each participant from their resting-state functional MRI correlation data, using the approach described in (Osher et al., 2019). This method is a variant of the method used in (Osher et al., 2016; Saygin et al., 2012, 2016), but applied to resting-state data. Briefly, we used the Glasser multi-modal parcellations from the Human Connectome Project(Glasser et al., 2016) to define a target region (within the fusiform) and search regions (the remaining parcels). The fusiform target region in each hemisphere included the following 5 Glasser parcels in each hemisphere – Area V8, Fusiform Face Complex (FFC), Area TF, Area PHT and Ventral Visual Complex (VVC). Next, we defined the “correlation fingerprint” for each vertex in the fusiform target region. The CF of each vertex was a 355-dimensional vector, each dimension representing the Pearson correlation between the time course of that vertex at rest, and the time course of one of the 355 remaining Glasser parcels in the search space during the same resting scan (averaged across all vertices in the chosen target Glasser parcel).

#### Predictive model

We next trained a ridge-regression model to predict the face-selectivity of each voxel in the fusiform target region directly from the correlation fingerprints(Osher et al., 2019). The T-statistic map from the contrast of face > maze, hand and chair were used for the predictions as in previous studies(Osher et al., 2019, 2016; Saygin et al., 2012). The model was trained using standardized data (mean centered across all vertices in the fusiform search space and unit standard deviation) using a leave-one-subject-out cross-validation routine. This method ensures that the individual subject data being predicted is never used in the training procedure. The ridge-regression model includes a regularization parameter which was determined using an inner-loop leave-one-subject cross-validation. Each inner-loop was repeated 100 times, each with a different coefficient (lambda values logarithmically spaced between 10^-5^ and 10^2^). The optimal lambda and betas were chosen from the inner-loop models that minimized the mean-square error between the predicted and observed activation values and used to predict the responses for the held-out subject. The model prediction accuracy was estimated as the Pearson correlation between the predicted and observed patterns of face selectivity across vertices in the fusiform target regions for each subject. We evaluated the predictions from the model against the benchmark of predictions from a random-effects group analysis using the general linear model (in Freesurfer) applied to the face-selectivity of the other subjects.

The model training and benchmark procedures were similar to (Osher et al., 2019), except for one important difference. We split the all of the functional and resting state data into 2 independent groups (even-odd run split). We trained all our models based on data from even runs and evaluated the model performances based on data from the odd runs. This additional procedure ensures that the predictions for each subject are independent samples for subsequent statistical analyses. Statistical significance was assessed using two-tailed paired *t*-tests on Fisher-transformed prediction scores across subjects. To assess the degree to which we could predict functional activations from parcels in individual lobes, we divided the search regions into frontal, parietal, occipital and temporal lobes based on the Glasser parcels(Glasser et al., 2016). These regions included 162, 66, 60 and 67 parcels each in the frontal, parietal, occipital and temporal regions respectively. We re-trained our models from scratch limiting the predictors to the parcels within the each of the lobes and evaluated the model performance and performed statistical tests as before.

## Supplementary Figures and Table

**Supplementary Figure 1:**
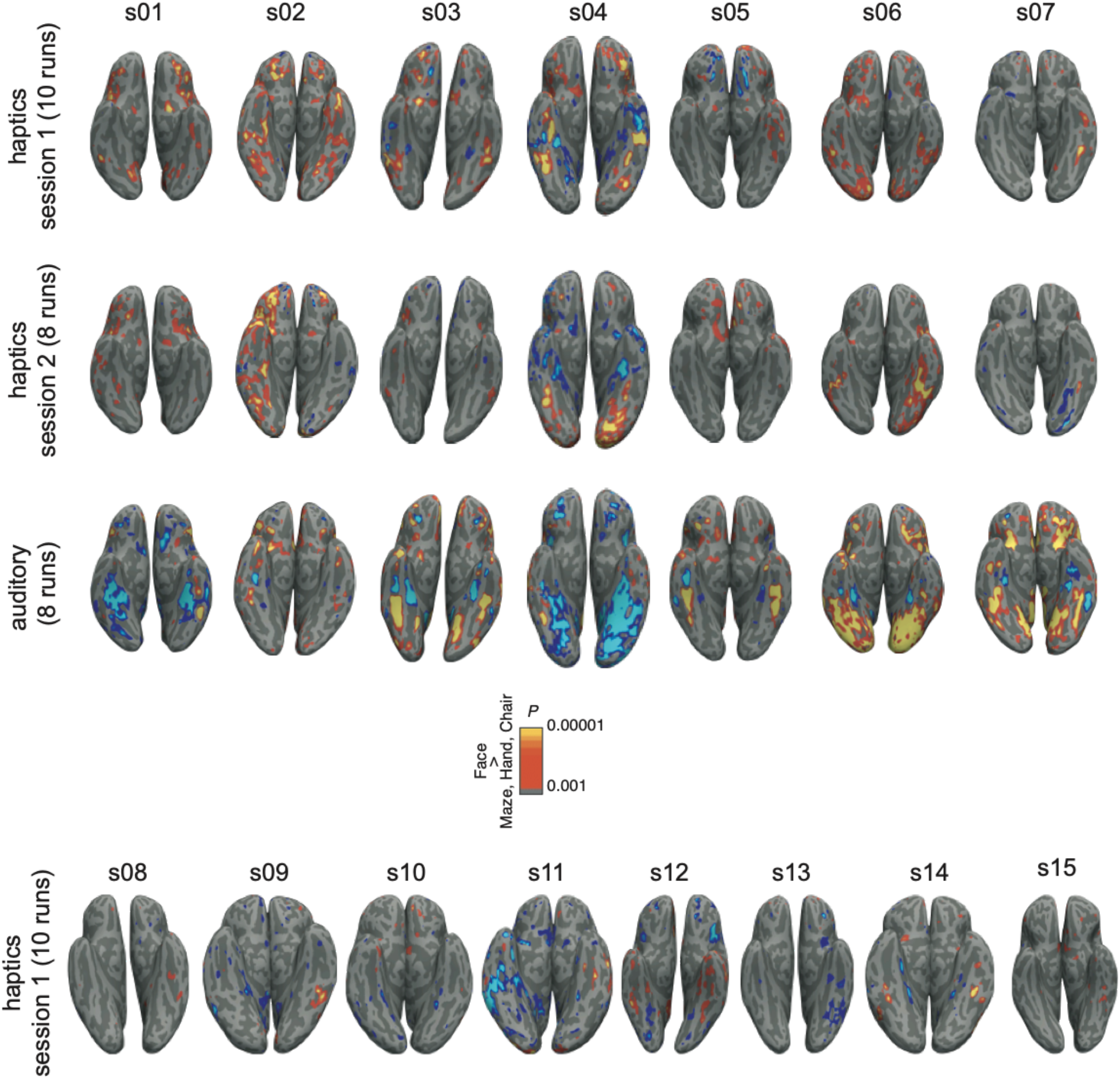
Activations in individual blind participants. Inferior view of each subjects’ native inflated surface reconstruction showing face-selective activations (Face> Body, Scene and Object, P<0.001, uncorrected). Subjects s01 to s07 (above the colorbar) performed all 3 Experiments. The top row shows the face-selective activations for Experiment 1 (haptics), the middle row for Experiment 2 (haptics with additional stimuli-spheroids and cuboids, although these conditions were not included in the face selectivity statistics shown here) and bottom row for Experiment 3 (auditory).

**Supplementary Figure 2:**
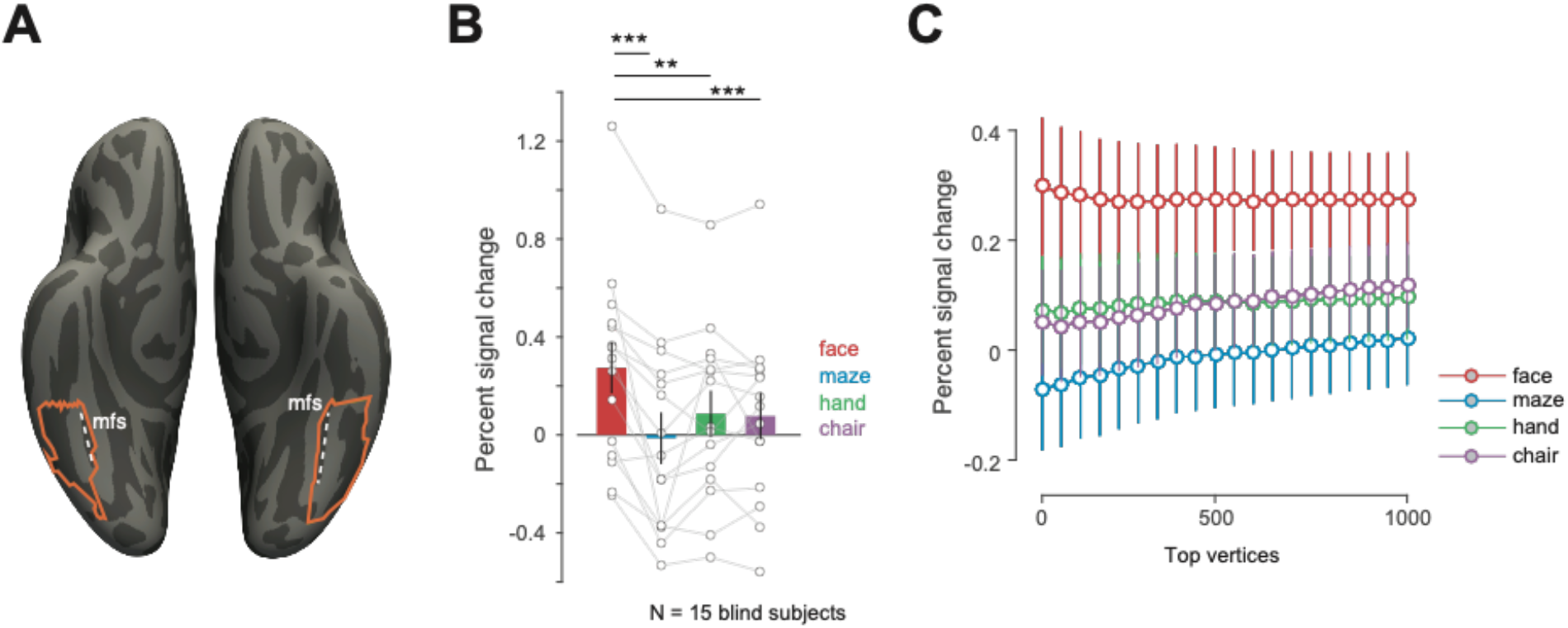
A cytoarchitecture-based method to identify face selectivity in the fusiform. **A.** An alternate anatomical constraint region within which to define face-selective activations in the fusiform, hand-drawn based on Weiner at al (2014) on the fsaverage surface. This parcel extends from the lateral side of the mid-fusiform sulcus (mfs) posteriorly to the posterior end of the collateral sulcus (ptCoS) **B.** Mean and s.e.m. of the fMRI BOLD response across blind subjects for the top 150 surface vertices (identified using independent data) during haptic exploration of 3D printed face, maze, hand and chair stimuli. ** is P<0.005 and *** is P<0.0005 **C.** Response profile of visual face-selective region (in held-out data) in blind subjects as a function of fROI size within the cytoarchitectonic-based anatomical constrain region in a (error bars indicate s.e.m.). The difference between face and object response magnitudes are statistically significant (P<0.005, two-tailed paired t-test across subjects) at each fROI volume up to 1000 vertices.

**Supplementary Figure 3:**
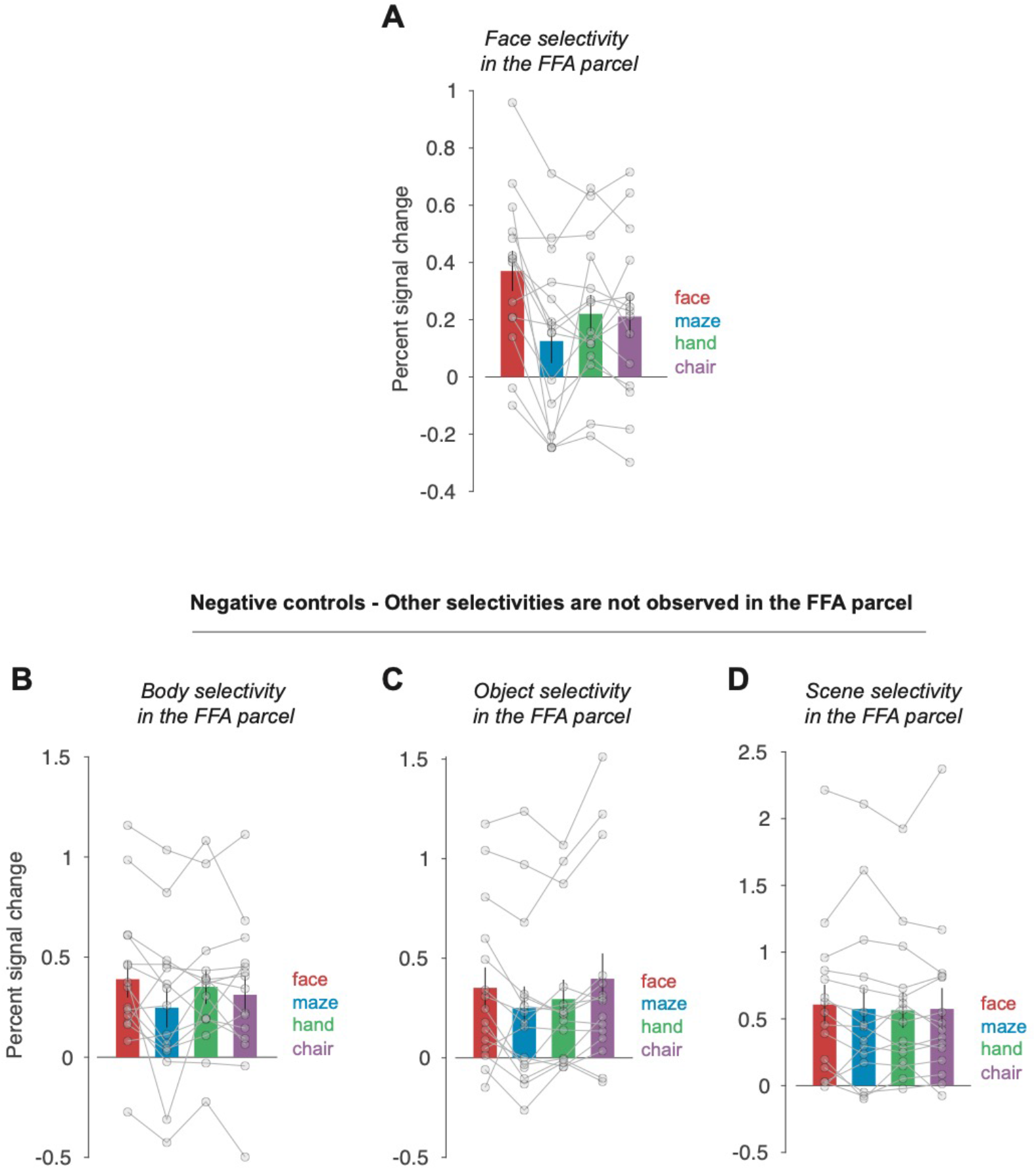
Negative controls for body, object and scene activations in group-constrained face parcels. **A.** Mean and s.e.m. for the BOLD response across blind subjects in the top 50 face-selective voxels (identified in independent data) during haptic exploration of face, maze, hand and chair stimuli. Individual subject data are overlaid as connected lines (same format as Figure 1K). **B,C,D.** Same as **A**, but for hand, chair and maze selectivity inside the face parcel. Note that none of these comparisons are statistically significant (all P>0.05). This analysis indicates that our analysis method does not produce false positives, detecting selectivities that should not exist in that region. Instead, the face parcels evidently constrain the location to identify face selective regions only and are not large enough to include regions selective for other object categories.

**Supplementary Figure 4:**
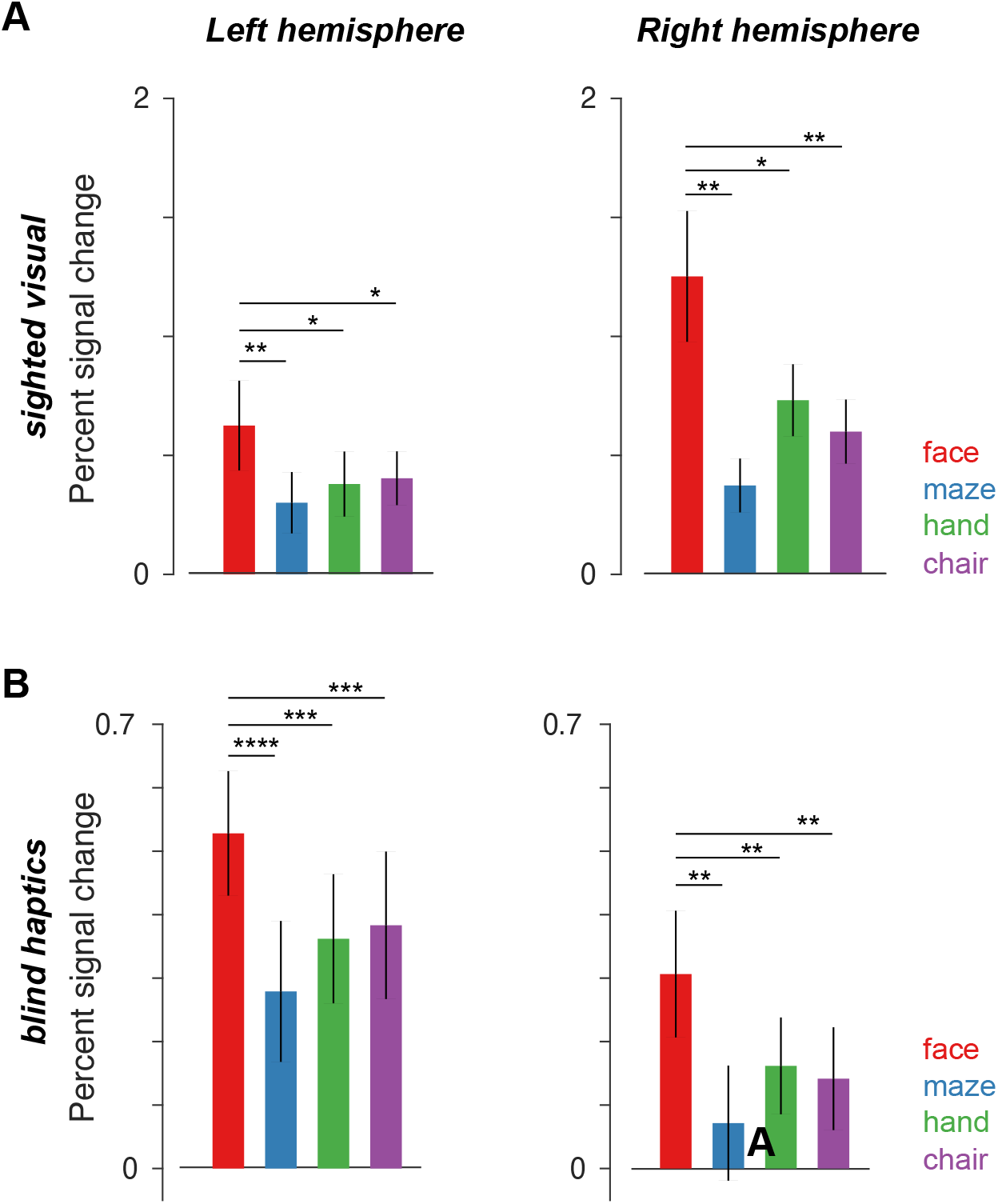
Lateralization of face selective activations in sighted and blind subjects. **A.** Mean and s.e.m. for the BOLD response across sighted participants (N = 15) in the top 20 face-selective voxels (identified in independent data) in the left and right hemispheres (analyzed separately) during visual exploration of face, maze, hand and chair stimuli. * is P<0.05, ** is P<0.005, *** is P<0.0005 and **** is P<0.00005 **B.** Same as A. but in blind participants (N =15) during haptic exploration of face, maze, hand and chair stimuli. A 3-way ANOVA on subject group (blind vs sighted), stimulus types (face, maze, hand and chair) and hemispheres (left and right) reveals a group by hemisphere interaction effect (F(1,227)=15.71, P=0.0001), indicating higher overall responses in the RH in sighted but in the LH in blind. However, there was no evidence of laterality differences between groups in selectivity: a) the triple interaction of group x hemisphere x stimulus condition was not close to significant (F(3,224) = 0.8, P=0.50), and b) a two-way ANOVA on selectivity index found no significant interaction of subject group by hemisphere (F(1,56) = 0.27, P=0.60) (and no main effect of either subject group, F(1,56) = 1.65, P=0.20) or hemisphere F(1,56) = 2.44, P=0.12)).

**Supplementary Table 1:**
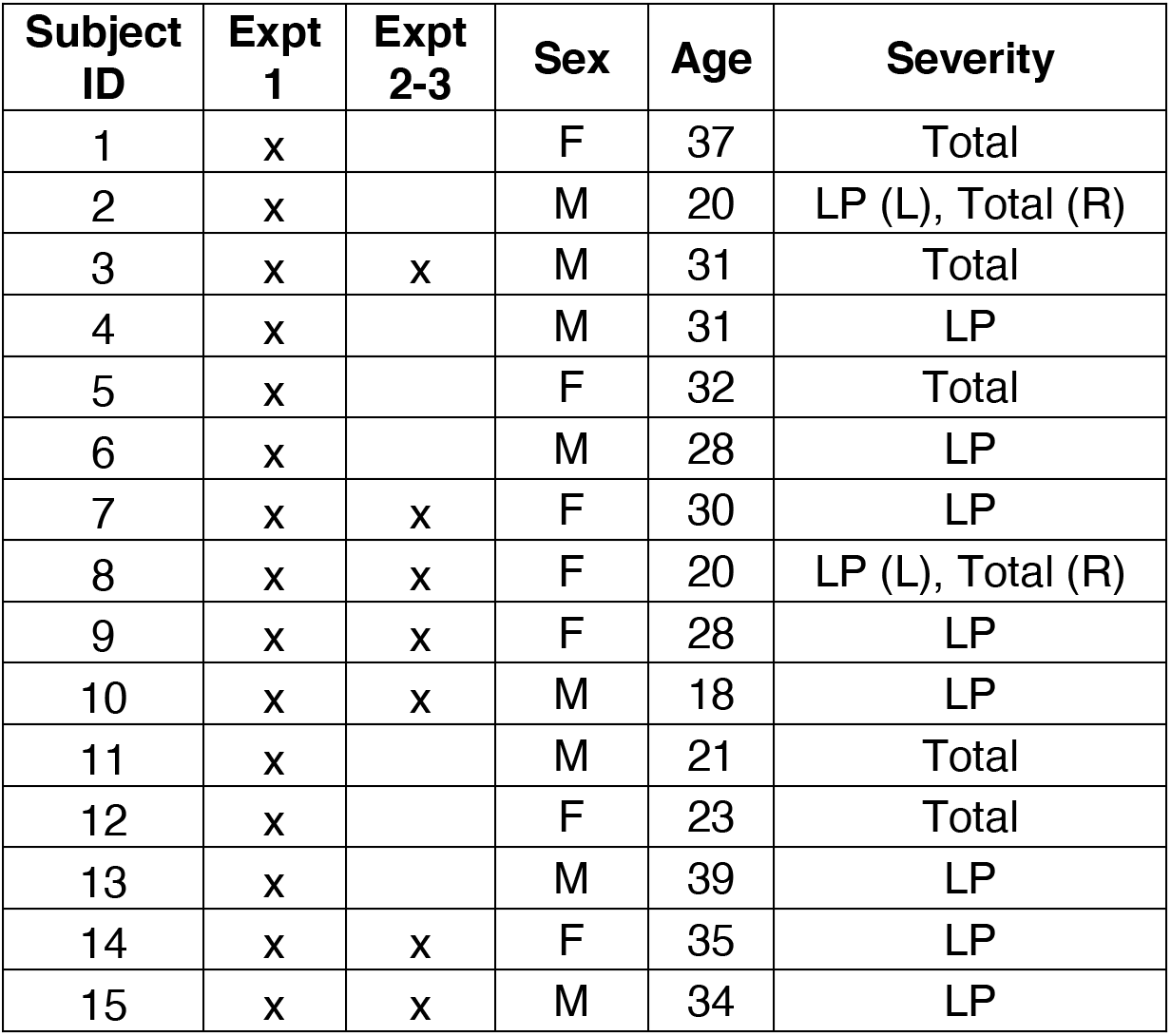
Blind participant details. Severity indicates level of blindness (for both eyes unless otherwise indicated). All recruited participants were congenitally blind (age of reported onset = 0y).

